# Following the footprints of variability during filopodia growth

**DOI:** 10.1101/2020.10.22.349084

**Authors:** Daniela Senra, Alejandra Páez, Geraldine Gueron, Luciana Bruno, Nara Guisoni

**Affiliations:** Instituto de Investigaciones Fisicoquímicas Teóricas y Aplicadas (INIFTA), Universidad Nacional de La Plata (UNLP), Argentina. Consejo Nacional de Investigaciones Científicas y Técnicas (CONICET), Argentina; Laboratorio de Inflamación y Cáncer, Departamento de Química Biológica, Facultad de Ciencias Exactas y Naturales, Universidad de Buenos Aires, Buenos Aires C1428EGA, Argentina.; CONICET-Universidad de Buenos Aires, Instituto de Química Biológica de la Facultad de Ciencias Exactas y Naturales (IQUIBICEN), Buenos Aires C1428EGA, Argentina.; Unidad de Transferencia Genética, Instituto de Oncología “Ángel H. Roffo”, Universidad de Buenos Aires, Buenos Aires, Argentina.; Instituto de Cálculo, CONICET-FCEN, Facultad de Ciencias Exactas y Naturales, Universidad de Buenos Aires, Pabellón 1, Ciudad Universitaria (1428), Buenos Aires, Argentina. Consejo Nacional de Investigaciones Científicas y Técnicas, Argentina.

**Keywords:** filopodia growth, stochastic model, actin, regulatory proteins, fluorescence microscopy, prostate cancer cells

## Abstract

Filopodia are actin-built finger-like dynamic structures that protrude from the cell cortex. These structures can sense the environment and play key roles in migration and cell-cell interactions. The growth-retraction cycle of filopodia is a complex process exquisitely regulated by intra- and extra-cellular cues, whose nature remains elusive. Filopodia present wide variation in length, lifetime and growth rate. Here, we investigate the features of filopodia patterns in fixed prostate cancer cells by confocal microscopy. Analysis of almost a thousand filopodia suggests the presence of two different populations: one characterized by a narrow distribution of lengths and the other with a much more variable pattern with very long filopodia. We explore a stochastic model of filopodia growth which takes into account diffusion and reactions involving actin and the regulatory proteins formin and capping, and retrograde flow. Interestingly, we found an inverse dependence between the filopodial length and the retrograde velocity. This result led us to propose that variations in the retrograde velocity could explain the experimental lengths observed for these tumor cells. In this sense, one population involves a wider range of retrograde velocities than the other population, and also includes low values of this velocity. It has been hypothesized that cells would be able to regulate retrograde flow as a mechanism to control filopodia length. Thus, we propound that the experimental filopodia pattern is the result of differential retrograde velocities originated from heterogeneous signaling due to cell-substrate interactions or prior cell-cell contacts.

## 1 Introduction

Filopodia are filamentous cell projections that protrude from the plasma membrane by the polymerization of actin filaments. Filopodia are well conserved structures, present in diverse cell systems and known to play a key role in cell migration, sensing and cell-cell communication [1, 2]. Typically, a filopodium contains a bundle of around 10-30 actin filaments [3] and grows at a speed of 0.01-0.2 µm/s [4], reaching lengths of a few micrometers [5–7]. As a consequence of the complex processes involved in filopodia dynamics there are wide variations in filopodial lifetimes, ranging from a few seconds to several minutes [8].

The mechanisms of filopodia initiation are still not clear. It has been proposed that filopodia would emerge as a consequence of the reorganization of the lamellipodial actin network upon the convergent elongation of privileged filaments, which would bind a complex of molecules to their barbed ends allowing them to continue elongating together [9]. On the contrary, other authors [4, 10] found that filopodia are able to form in the absence of lamellipodia suggesting different functionality between both actin structures.

Once initiated, the bundle of actin filaments elongates by the polymerization of actin monomers at the filaments barbed ends and retracts as a consequence of both, depolymerization and retrograde flow. Filopodia extension and retraction is a dynamic process controlled by different proteins, including capping protein and formin [11]. These regulators have opposite effects on the filopodia length: while formin accelerates actin assembly, capping protein prevents its polymerization [12, 13]. Both proteins bind actin barbed ends with high affinity and slow dissociation, which initially led to the conclusion that they were mutually exclusive [11, 14]. However, it has been recently shown that formin and capping proteins are able to simultaneously bind barbed ends and coregulate filament assembly [11].

It has been suggested that retrograde flow in the cell cortex at the base of the filopodium is the main retraction mechanism, which also generates retrograde forces on the substrate [15]. Retrograde flow originates from the combination of two processes: the actin treadmilling due to the depolymerization at the rear part of the filopodium and the action of myosin motors [16, 17]. For example, it has been found that filopodium elongated more that 80% after inhibition of myosin II in neuronal growth cones [18]. Depolymerization of actin at the tip can also contribute to the retrograde flow in melanoma cells and fibroblasts in a process regulated by cofilin and fascin [19].

The magnitude of the retrograde flow depends on the cell type and species; values in the range 5-260 nm/s have been reported (see references in Table 1 in [3]). Furthermore, its value might also depend on the state of development of the filopodia [20], on probes acting as guidance cues to their growth stabilizing or destabilizing filopodia [21–24] and/or on the substrate stiffness [25].

**Table 1.**
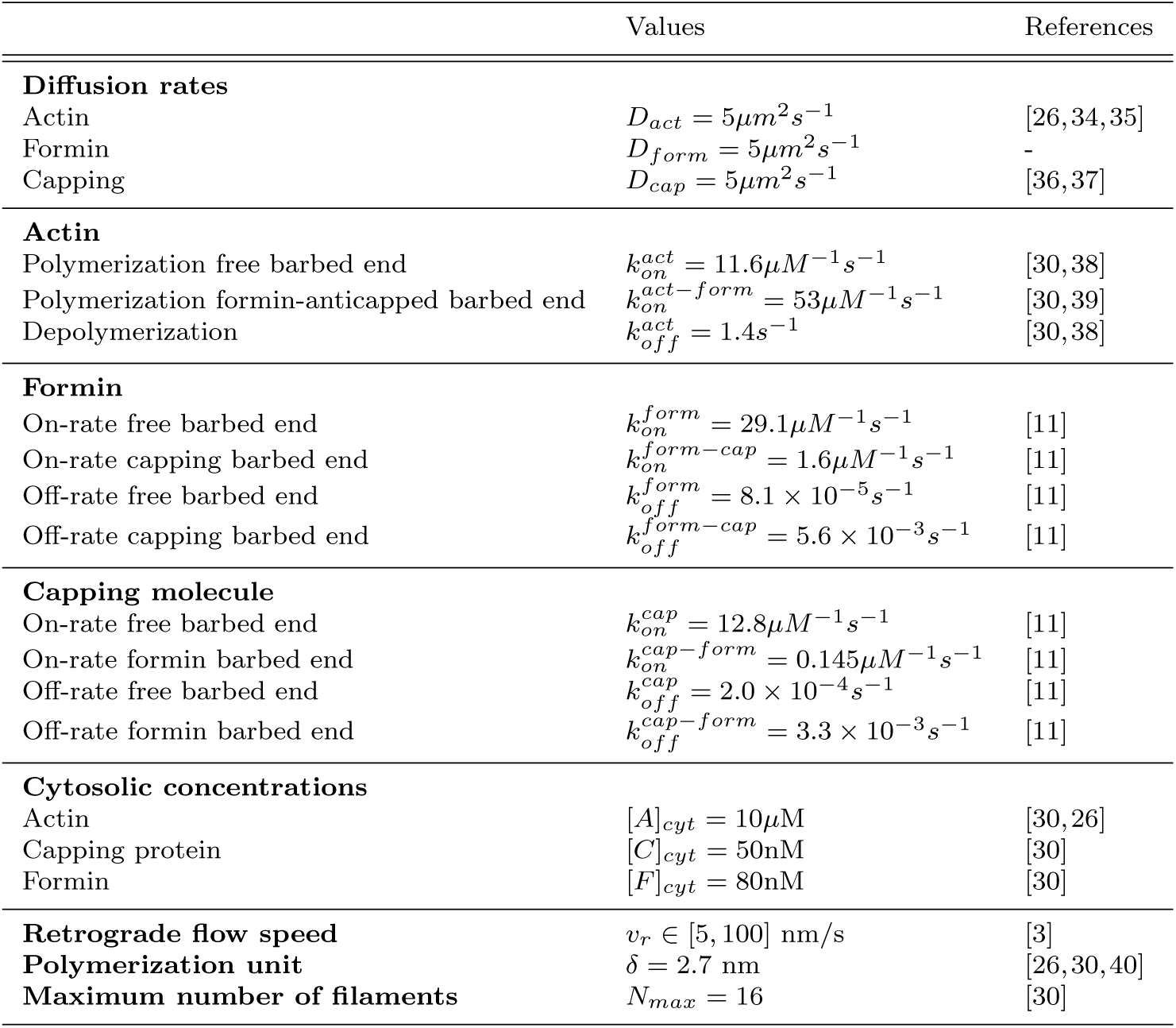
Parameters used in the model simulations.

The complexity of the mechanisms underlying the initiation, maintenance, and retraction of filopodia has inspired theoretical and computational approaches for a better understanding of the biological system. Regarding filopodia growth, several authors have addressed this issue, using deterministic analytical models [7, 26–28] and stochastic simulations [29–32]. In a groundbreaking publication, Mogilner and Rubinstein [26] propose a deterministic one-dimensional reaction-diffusion model for G-actin dynamics within filopodia. Some characteristic scales of filopodia emerge from the model, such as the typical filopodia length (of the order of few microns) and that more than 10 bundled filaments are required to avoid buckling. Other authors propose mean-field models that also account for the effect of myosin motors [7, 28] and capping proteins [27]. On the other hand, Papoian’s group has made a significant contribution to the stochastic modeling of filopodial growth [29–31, 33]. In this way, Lan et al. [29] developed a one dimensional model for a bundle of filaments within a filopodium. They consider the polymerization and depolymerization of actin at the barbed end, as well as an effective retrograde velocity for each filament individually. The model allows to observe an equilibrium state with fluctuations around the stationary length. Zhuravlev and Papoian [30] brings complexity to this model, by adding the regulatory effect of capping proteins and formins. Interestingly, considering actin-binding proteins amplifies molecular noise and eventually leads to large growth-retraction oscillations in the filopodial lengths.

In this paper, we study the pattern distribution of filopodial lengths by means of a stochastic model based in the work of Zhuravlev and Papoian [30], though considering recent results showing that formin and capping proteins coregulate the actin assembly [11]. It has been hypothesized that cells can control different processes such as migration or the establishment of cell-cell contacts by regulating the retrograde flow and, as a consequence, the length and stability of filopodia [21, 23, 25]. Thus, we explore *in silico* the effect of varying the magnitude of the retrograde flow on the mean filopodial length and, more importantly, on its dispersion. We compare the results of the numerical simulations with the filopodial lengths obtained from the inspection of filopodia in fixed cultured prostate cancer cells. Our results suggest that the experimental data would consist of at least two populations of filopodia characterized by different retrograde velocities. The biological significance of our results is also discussed.

## 2 Results

### 2.1 Stochastic model of filopodia growth

We study filopodia growth dynamics by considering a stochastic reaction-diffusion model. The model used here is similar to the proposed by Zhuravlev and Papoian [30]. A growing filopodium generates a region that is an extension of the cytoplasm where actin, formin and capping molecules can react and diffuse (see Fig. 1a). By using a molecular approach [31] the chemical reactions at the filament barbed end are explicitly considered. The depolymerization at the filament pointed end as well as the action of the myosin motors are considered as an effective process that regulates filopodia retraction. The diffusion of the distinct molecules into the cytoplasmic extension is taken into account explicitly.

**Fig. 1.**
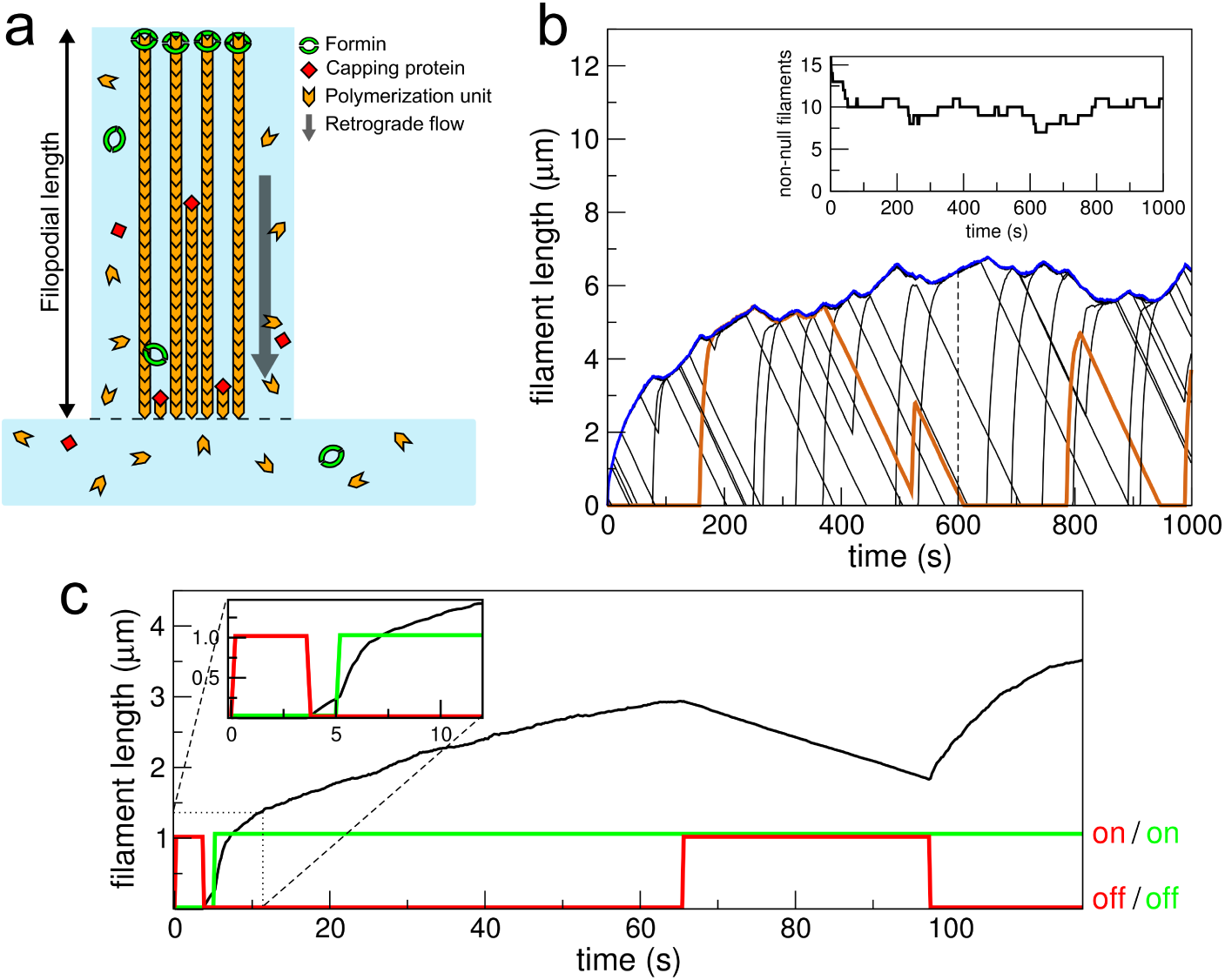
(a) Schematic representation of the model for filopodial growth. (b) Filaments within a filopodium length and filaments with non-null length (inset) as a function of time for *v*_*r*_ = 30 nm/s. The filopodium length corresponds to the largest filament (blue line). As an example, for 600 s (dashed line) there are two short filaments, one with intermediate length, and several ones with maximum length (schematically illustrated in (a)). The orange line highlights the dynamics of a single filament, showing that it can disappear and grow again from the cytoplasm. (c) Filament length and binding/unbinding of formin and capping molecules (black, green and red, respectively), as a function of time for *v*_*r*_ = 10 nm/s. Formin increases the growing filament velocity (~ 5 s) whereas capping blocks filament polymerization, and the length decreases due to the retrograde flow. In the time interval between 65 and 97 s both formin and capping are bound. Inset: enlargement of short time region.

Following the model proposed by Zhuravlev and Papoian [30], the filopodia structure is built with actin molecules as building blocks: G-actin polymerizes into F-actin at the filament barbed end. Actin depolymerization of the filament barbed end is also possible. These processes occur with rates 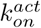 and 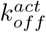, respectively. Formin and capping proteins can also bind at the filament barbed end, noticeably affecting G-actin polymerization. In this way, while capping of the filaments blocks actin polymerization, formin enhances substantially filament assembly [12, 13]. Therefore, if 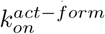 is the actin polymerization rate when a formin molecule is already bound to the filament, we consider that 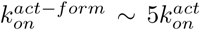 [39]. In our model we also considered recent results showing that formin and capping proteins can coregulate filament assembly by simultaneous binding to the barbed end [11]. This aspect was not taken into account by Zhuravlev and Papoian [30], who assumed that formin and capping were mutually exclusive. However, Shekhar and coauthors [11] found that the on- and off-rates of formin (capping) are affected by the presence of the capping (formin) protein [11]. We allow for these results by defining new values for the binding and unbinding rates 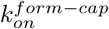 and 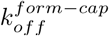 binding of formin when a capping molecule is already bound to the filament, and 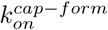 and 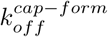 for the corresponding capping case.

G-actin, formin and capping molecules cytosolic concentrations are considered to be constant, with bulk values [*A*]_*cyt*_, [*F*]_*cyt*_ and [*C*]_*cyt*_, respectively [30, 26]. These molecules are able to diffuse into the cytoplasmic extension with effective diffusion coefficients *D*_*act*_, *D*_*form*_ and *D*_*cap*_ [26, 34–37]. This is another difference in relation to the work of Zhuravlev and Papoian [30], which does not take into account the diffusion of formin and capping molecules. Formin diffusion rate was considered with the same value as actin since both have similar molecular weight. Finally, the depolymerization of actin at the pointed end coupled with the action of myosins are taken into account as an effective process that regulates filopodia retraction. In this way, we assume a constant retrograde velocity whose action is to continually shorten the filaments, as made previously [29–31].

Despite that an actin filament is composed of two protofilaments [41], we model each filament as a rodlike structure without contemplating its internal double stranded helical organization, as it was also considered in previous works [26, 29–31, 33, 40]. The successive binding of actin molecules makes the filament length increase by a polymerization unit, as schematically shown in Fig. 1a. Also, we consider that a bundle of filaments constitutes a filopodia, with a maximum number of 16. The filopodia length is equal to the largest filament length (Figs.1a and b). We use a stochastic molecular approach to simulate reactions and diffusion in our model [31], as discussed in Section 4.1. The parameters used in the model are given in Table 1.

A representative filament temporal evolution is shown in Fig. 1c. The growth velocity of the filament is notably increased by the binding of a formin, as can be seen in the inset of Fig. 1c. On the other hand, when a capping molecule binds to the filament, polymerization stops and the filament retracts linearly as a consequence of the retrograde velocity.

Fig. 1b shows an example of the dynamic of the filaments within a filopodium. The number of filaments with non-null length is shown in the inset. According to the model, the filopodia growth rate is high for short times, but as time goes on, the velocity decreases exponentially. This result is in agreement with the deterministic model proposed by Mogilner and Rubinstein [26]. In fact, for longer times a balance between the actin polymerization and the retrograde velocity leads to a stationary length. This result is in accordance with the fact that the number of non-null filaments tends to an average value (inset of Fig. 1b). As can be seen in Fig. 1b, if one filament disappears by the continuous binding of a capping molecule it is able to grow again from the cytoplasm. Notably, the stochastic binding/unbinding of formin and capping molecules results in high variability in filopodial length (also observed in [30]), as compared to models where only actin polymerization/depolymerization is taken into account [29, 31].

The filopodial length is obviously affected by the model parameters. An elegant mean field expression for the steady-state filopodium length was obtained for models where only actin polymerization/depolymerization is considered [29, 33]. These authors found that the stationary length increases linearly with both the diffusion coefficient of actin and the actin cytosolic concentration. Their results also showed that the steady-state filopodium length presents a nonlinear increase with the binding rate of actin and a nonlinear decrease with both the retrograde velocity and the number of filaments. Finally, the stationary filopodium length is not significantly influenced by actin depolymerization. Here we are interested in studying the effect of varying the model parameters that may be involved in the regulation of filopodia growth. We assume that biochemical processes intrinsic to the cell, such as polymerization/depolymerization rates, diffusion coefficients, cytosolic concentrations, and the number of filaments in a filopodium, will not be highly variable among cells with a common identity. The sensitive nonlinear behavior of the stationary filopodium length with the retrograde velocity observed for simpler models suggests that retrograde flow can work as a control mechanism of filopodia growth and retraction [29]. Therefore, we will explore the effect of varying the retrograde velocity throughout the simulations.

#### 2.1.1 Influence of retrograde velocity in filopodia length

In order to explore the effect of the retrograde flow on the filopodial lengths, we run simulations varying the retrograde velocity (*v*_*r*_) while keeping the rest of the parameter values unchanged (Table 1). Fig. 2a displays the results for 3 different values of *v*_*r*_ in the range between 10 and 100 nm/s. For each value of *v*_*r*_, Fig. 2a presents several individual simulations, as well as the filopodial length averaged over these curves (for a definition see Eq. 3 in section 4.2). Our simulations go up to *t* = 1000 s, since the average filopodia lifetime is reported to be in the order of a few minutes [5, 42, 43], even though some filopodia are longer-lived [42–44]. Interestingly, the time course of individual filopodia lengths shows large fluctuations, and different simulations for the same parameter values are also very variable. The reason for such behaviour lies on the stochastic nature of the binding/unbinding of actin and regulatory proteins, taken into account by the model. We verified that larger values of *v*_*r*_ resulted in shorter lengths. It can also be observed that for low values of *v*_*r*_, the filopodial length presents a larger transient towards the stationary length. Also, the filopodium growth rate (estimated from the average speed at each simulation time step in the short time region, see Fig. 2a) is of the order of 100 nm/s, comparable with the reported values [4].

**Fig. 2.**
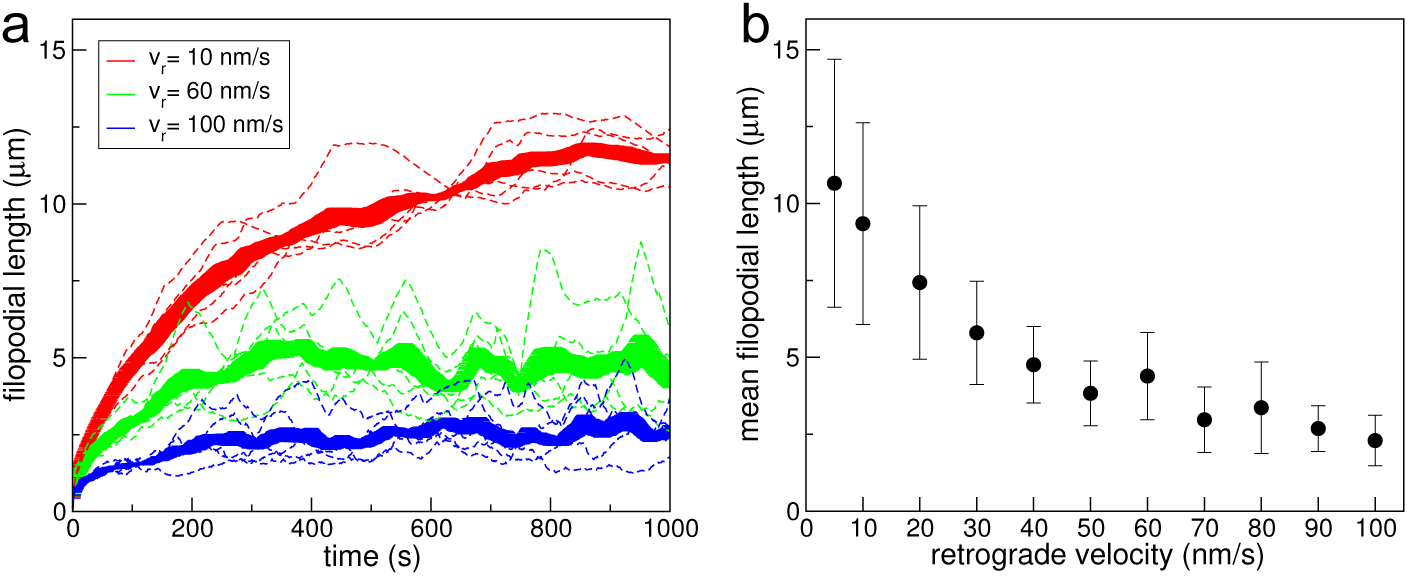
(a) Filopodial length (dashed lines) for different retrograde velocities *v*_*r*_, as indicated. For each value of *v*_*r*_ the solid line represents the filopodial length averaged over the 5 runs, 〈*L*(*t*)〉, and the thickness of the line depicts the standard error. (b) Mean filopodial length 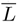 as a function of *v*_*r*_. The error bars represent the standard deviation. Further information about simulation data can be found in Section 4.2.

Many experiments deal with images of fixed cells [5, 6, 45, 46], which display the distribution of filopodia at arbitrary moments in their lifetimes. To take this type of data into account, we will consider the full-time behaviour of filopodial length given by the model, instead of only the stationary value, as it was previously done in other theoretical works [29]. Therefore, we compute the average of the filopodial lengths over both time and different realizations as an estimate of the mean filopodia length (Eq. 4 in Section 4.2). Fig. 2b shows that the mean filopodial length exhibits a nonlinear inverse dependence on the retrograde velocity. Then, larger values of *v*_*r*_ result in shorter filopodia. Furthermore, it can be seen that the mean filopodial length is much more sensitive to variations of *v*_*r*_ for low values of retrograde velocity than for higher ones. The large dispersion in the mean filopodial length observed in Fig. 2b for low values of *v*_*r*_ is associated with the existence of a long transient to reach the equilibrium length (see Fig. 2a).

### 2.2 Filopodia in PC3 cells

In order to explore filopodia patterns, we studied cultured prostate cancer cells (PC3 cells). After being fixated and stained for actin with rhodamine-phalloidin, cells were imaged by confocal microscopy [45]. While this condition has the advantage of allowing inspection of all the cells in the field with little noise, it has the drawback that the dynamical behavior of filopodia cannot be explored. However, these images are representative of the distribution of cells and filopodia in an arbitrary moment of their cycle. The images were analyzed with custom made routines. A detailed description of the image analysis can be found in Section 4.4.

We show PC3 cultured cells imaged using phase contrast optics in Fig. 3a. The micrograph in Fig. 3b corresponds to the fluorescence channel and in Fig. 3c we exhibit an overlay of both images. Since filopodia are easily distinguished in the fluorescence images, we found it more convenient to use these images for our analysis.

**Fig. 3.**
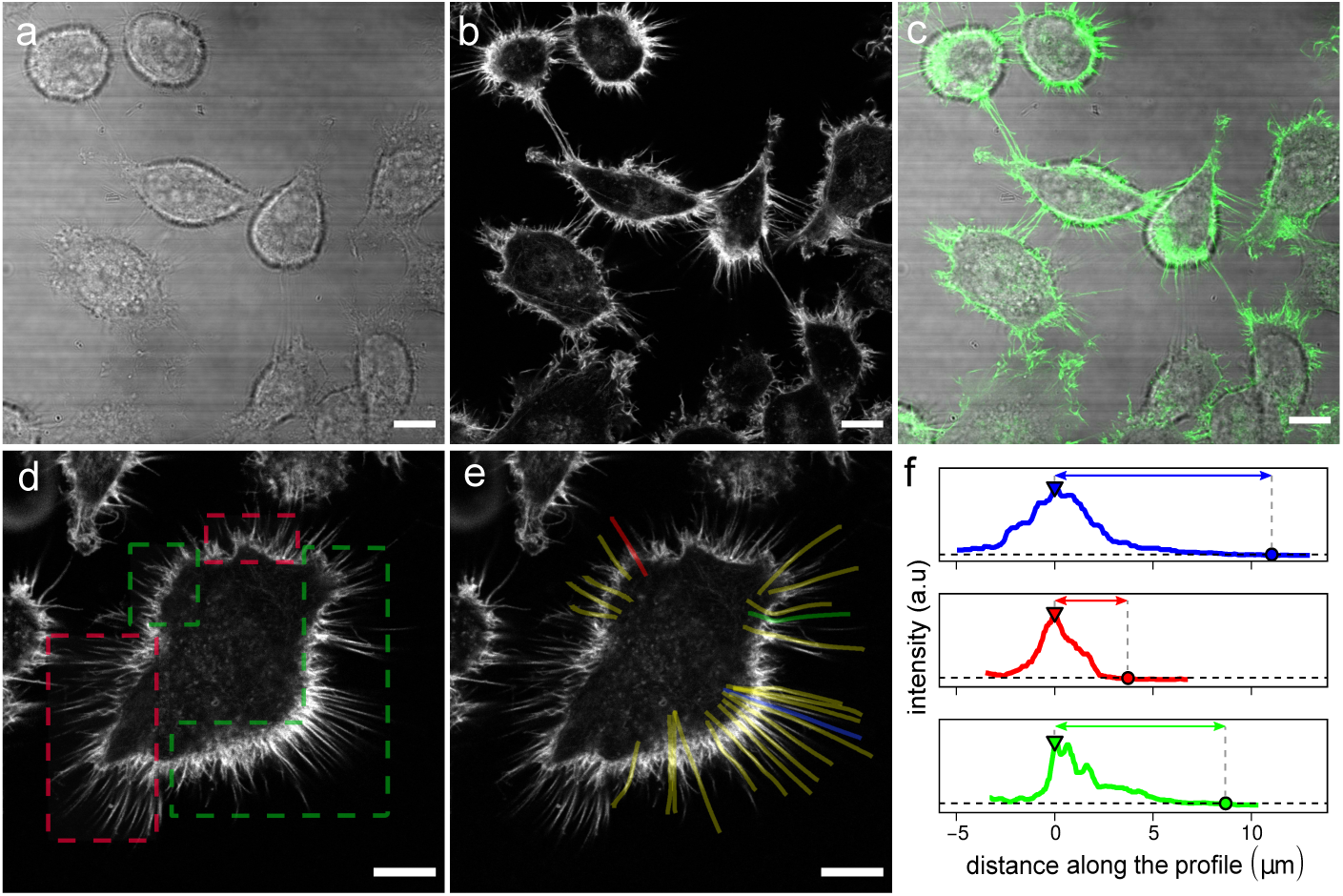
Prostate cancer PC3 cells. (a) Phase contrast microscopy image. (b) Fluorescence microscopy image of the same viewfield (staining with rhodamine-phalloidin). (c) Merge of the fluorescence and phase contrast confocal images shown in a and b. (d) Representative fluorescence image of PC3 cells. The regions of interest where filopodia are considered trackable are delimited by green shapes. The red rectangles surround regions where filopodia are considered unfit to be measured. In this case, the upper-right red rectangle encloses a contact region and the lower-left shape too, the latter also includes overlapping protrusions with intermittent intensity signal. (e) Lines indicate the filopodia that could be tracked (Scale bars: 10 *µm*). (f) Intensity smoothed profiles acquired from the lines traced in (e); the colours of the profiles correspond to the filopodia in (e). The filopodial tip locations are determined after selecting a threshold that considers background intensity (black dashed horizontal line). The triangles and circles represent the base and tip positions respectively. The protrusion length is obtained by subtracting tip and base positions along the profile. Filopodial length of the protrusions covered by the blue, red and green lines in e are represented by lines with arrowheads of the respective colour. The smoothed profiles are horizontally shifted so that the filopodial base is at the origin.

We focused on individual filopodia and not on filopodia located at cell-cell junctions or that form bridge-like structures, such as those described in [47]. Thus, we identified membrane regions where filopodia represented clearly delimited structures, while cell-cell protrusions and extremely dense regions were excluded for the analysis, similarly to [43]. Furthermore, filopodia with low signal to noise ratio (low intensity with respect to the background) and/or intermittent intensity levels that might correspond to protrusions that enter and go out the image plane were considered unfit to be measured. Fig. 3d shows an example where the discarded areas were delimited by red rectangles and the trackable regions were circumscribed by green shapes.

We sampled filopodia from the trackable zones, generally one or two regions per cell were explored. We identified individual filopodia from the fluorescence images and manually delineated them (Fig. 3e) to acquire intensity profiles (Fig. 3f). We input the collected intensity profiles into a custom made program which determines the filopodial tip and base locations for each profile and computes the filopodial length by substracking them. Thereby, we measured the length of approximately a thousand filopodia.

To compute the proportion of filopodia sampled in the explored regions, we measured the contour length of 45 sampled areas and multiplied it by the filopodial linear density obtained by Paez et al. [45] from the same images. We estimated that half the filopodia population within the inspected areas were roughly sampled. We summarized the image analysis statistics in Table 2.

**Table 2.** Image analysis statistics.

| Symbol | Meaning | Value |
| --- | --- | --- |
| $N_{im}$ | No. of analyzed images | 48 |
| $N_{cells}$ | No. of cells explored | 226 |
| $N_{filo}$ | No. of measured filopodial lengths | 994 |
| $\hat{p}$ | Estimated percentage of sampled filopodia | 56% |

### 2.3 Analysis of filopodial lengths in cultured PC3 cells suggest two cell populations

We analyzed more than a thousand filopodia profiles from which we recovered 994 filopodial lengths in the range 0.5 to 15 *µm*. The average filopodial length for cultured PC3 cells was ~ 4 *µm*, which is similar to the lengths of filopodia in human lung adenocarcinoma cells [25], rat fibroblasts [6] and neurites [5]. Fig. 4a displays the obtained distribution; a quantitative description of the distribution in terms of it central moments is found in Table 3. The distribution is biased towards small length values and is long-tailed to the right as both the skewness and excess kurtosis apart from zero. Similar distributions for filopodial lengths were reported in [5, 6].

**Table 3.** PC3 filopodial length statistics.

| Filopodia | $N_{filo}$ | Mean [$\mu m$] | std | Skewness | Excess kurtosis |
| --- | --- | --- | --- | --- | --- |
| all | 994 | 4.1 | 1.9 | 1.7 | 3.9 |
| >6 | 561 | 4.1 | 2.0 | 1.7 | 3.9 |
| group 1 | 319 | 3.4 | 1.1 | 0.6 | 0.6 |
| group 2 | 242 | 5.1 | 2.6 | 1.1 | 1.0 |

**Fig. 4.**
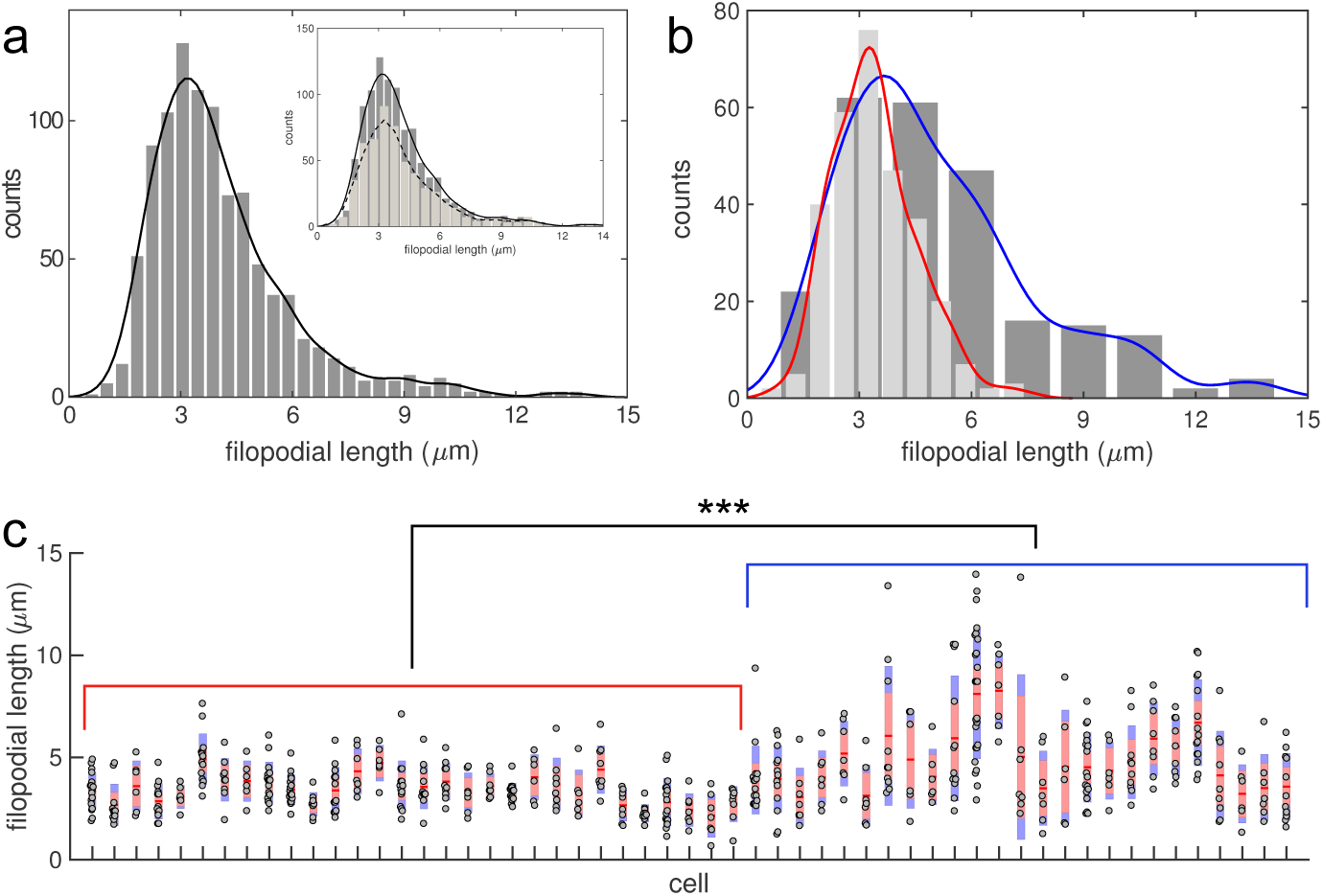
(a) Distribution of filopodial length in cultured PC3 cells. Inset: Comparison with the distribution of filopodial lengths sampled from cells where 6 or more filopodia could be tracked (light gray). (b) Histograms of the filopodial lengths corresponding to Group 1 (light gray) and Group 2 (dark gray) (see main text). (c) Distribution of single cell data corresponding to Group 1 (red brackets) and Group 2 (blue brackets). Boxplots are presented as mean, SEM and SD, according to [49]. Asterisks indicate significant differences between the two groups according to a two-sample Kolmogorov-Smirnov test. The lines over the histograms represent re-normalized kernel density estimates.

We then wondered if this wide distribution of filopodial lengths results from local variability or it stems from a variability between different cells. In order to explore this, we analyzed the lengths of neighbor filopodia located within the same cell. To this end, we gathered the data coming from cells where we were able to successfully track 6 or more filopodia profiles (*N*_*filo*_=561 from *N*_*cells*_=58) to allow meaningful analysis. The resulting data (inset in Fig. 4a was representative of the total data set as reflected by the distribution statistical description shown in Table 3.

As an estimate of the dispersion of the filopodial lengths within single cells, we computed the standard deviation (SD) for each of the 58 cells (not shown) and found that their distribution deviates from normal according to a Shapiro-Wilk hypothesis test (*p − value* < 5.10^−5^) [48]. Thus, we classified the cells into two groups depending whether the SD of the lengths was less (Group 1) or greater (Group 2) than SD=1.3 *µm*. This threshold value roughly corresponds to the median of the SDs distribution. The new distributions obtained for Group 1 (*N*_*filo*_=319) and Group 2 (*N*_*filo*_=242) are displayed in Fig. 4b, where we have also plotted the corresponding kernel density estimation. A quantitative description of the distributions in terms of their central moments can be found in Table 3. A two-sample Kolmogorov-Smirnov test showed that the two groups were significantly different (*p value* < 10^−14^) supporting the presence of at least two different population of data. While one group of cells (Group 2) displays a wide distribution of lengths with very long filopodia, the other group (Group 1) shows a narrow and less biased distribution, with the absence of long processes.

To assess whether our previous observation was correct, we went back to the images to determine if the filopodia tracked in cells coming from Group 1 were representative of the ensemble of filopodia in the corresponding group. In other words, we seek the presence of long filopodia within this group that would have not been considered during the tracking procedure. To this end, a line separating a distance *d* = 8 *µm* from the cell perimeter was drawn and the number of filopodia exceeding this limit was inspected. Also, we estimated the length of long curled filopodia within this 8*µm*-sized zone. This value of the length cutoff represented an upper bound for the Group 1 lengths, and thus filopodia longer than this value should represent outliers. This study revealed that only three of the considered cells displayed a least one filopodium longer than this threshold and, consequently they were discarded from further analysis.

Fig. 4c displays the distribution of filopodial lengths within the remaining 55 individual cells, where we have intentionally sorted cells depending on the group they belong to. An inspection of this figure also evidences the differences between the two populations of filopodia previously remarked.

### 2.4 Interpretation of the experimental data using the stochastic model

When comparing the experimental data for PC3 cells with the results of the stochastic model, it led us to formulate two questions. Can the stochastic model reproduce the experimental results for PC3 cells? What can we learn from that?

To answer to these questions, we first estimated the probability density functions of the filopodium lengths obtained from the stochastic model by performing a kernel density estimation. Some examples are shown in Fig. 5a, for five different values of *v*_*r*_. We called *E*_1_ and *E*_2_ the experimental probability density functions for Groups 1 and 2, respectively, which are obtained from a kernel density estimation of the experimental data (see Fig. 4b). Then we propose to reconstruct *E*_1_ and *E*_2_ as a linear combination of the simulated distributions *S*_*j*_, where the sub-index *j* stands for the *j − th* retrograde velocity value:

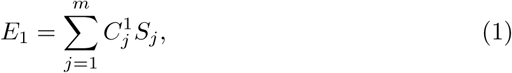

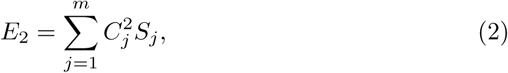

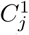 and 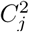 are the weight of the contribution of each theoretical distribution for Groups 1 and 2, respectively. We regard the same *m* = 11 values of retrograde velocity covering the range between 5 and 100 nm/s for both Eqns. 1 and 2.

**Fig. 5.**
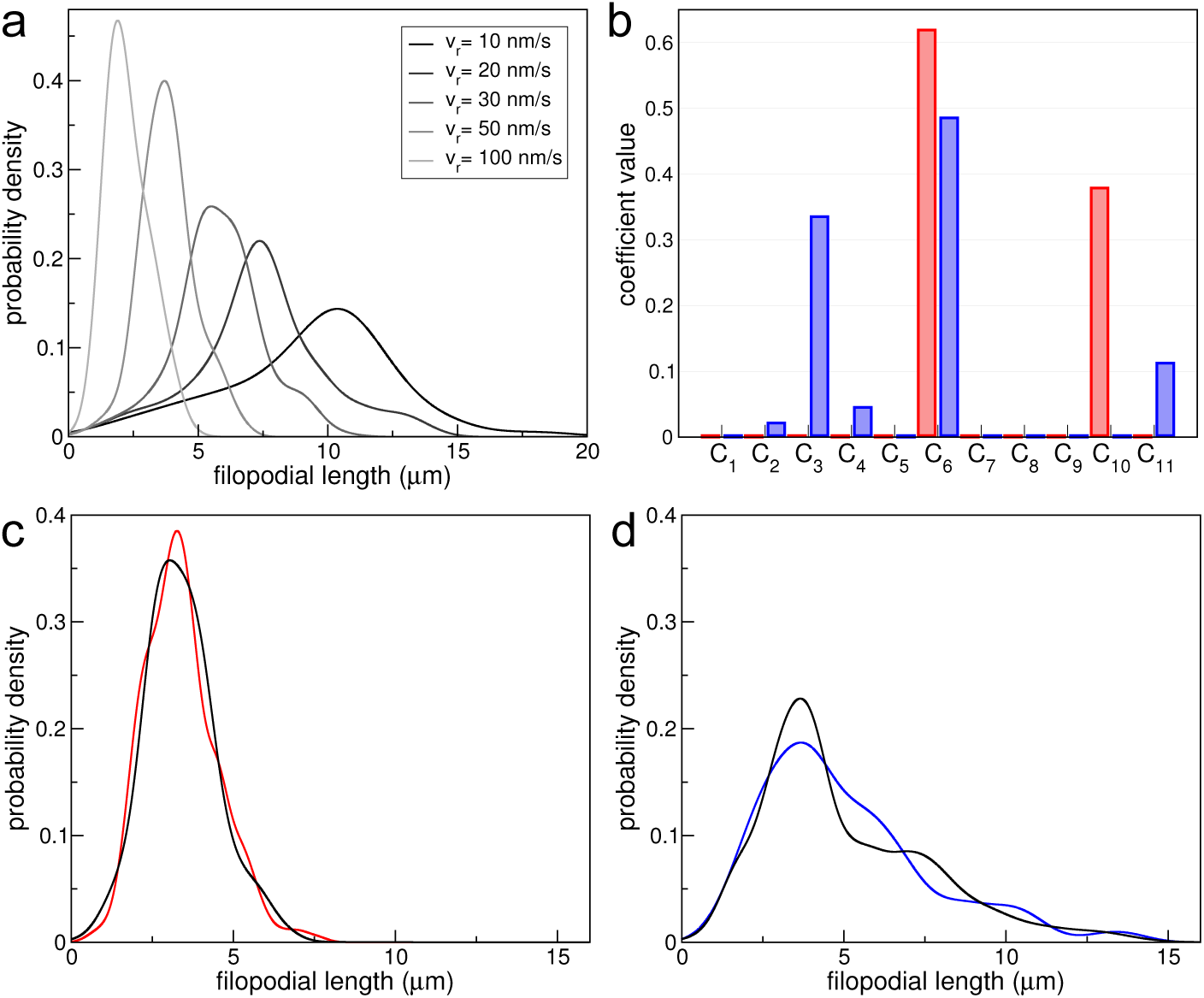
Comparison between experimental and simulation data using 4 moments of the distributions. (a) Probability density functions of the theoretical filopodium lengths for different values of retrograde velocity as indicated in the legend, obtained from kernel density estimations. (b) Coefficients of Eqns. 1 and 2 for Groups 1 and 2 (red and blue, respectively). For Group 1 the coefficients 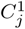 whose contribution is greater than 2% are related to *v*_*r*_= 50 and 90 nm/s (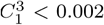, it is smaller than the line thickness). For Group 2 the non-null coefficients 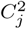 are associated to *v*_*r*_ = 10, 20, 30, 50 and 100 nm/s. (c) and (d) Experimental distributions *E*_1_ and *E*_2_ (red and blue lines, respectively), and the reconstructed simulation data by considering the coefficients shown in panel (b), for Groups 1 (c) and 2 (d).

We have shown in Table 3 that the Groups 1 and 2 skewness and excess kurtosis apart from zero. Therefore, we use the first four moments of *E*_1_, *E*_2_ and *S*_*j*_ to calculate the coefficients 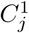 and 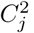, as discussed in Section 4.5. The coefficients obtained from Eqns.1 and 2 are displayed in Fig. 5b. Figs. 5c and d exhibit the reconstruction of *E*_1_ and *E*_2_ using these coefficients, which shows the good agreement between the model and the experimental data.

Note that contributions of *v*_*r*_ in the range 10 to 100 nm/s are necessary to reconstruct Group 2 distribution, while Group 1 distribution can be recovered with *v*_*r*_ values between 50 and 90 nm/s (by only considering contributions with weights greater than 2%). This result is related with the dispersion of the length values, which is greater in the case of Group 2 (see Table 3). Also, there is a predominance of low and medium values of *v*_*r*_ for Group 2, that is, *v*_*r*_ = 20 and 50 nm/s 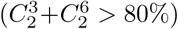. For Group 1, the emphasis is in medium and high values, that is, *v*_*r*_ = 50 and 90 nm/s 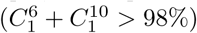. Since lower values of *v*_*r*_ are associated with higher values of filopodial length, the result above responds to the fact that Group 2 presents a long-tailed distribution biased to high length values.

Therefore, the main lesson from the comparison between the experimental data for PC3 cells and the stochastic model is that the reconstruction of Group 2 data involves a wide range of values of *v*_*r*_, and that the main contributions come from low *v*_*r*_ values.

## 3 Discussions

Filopodia play key roles in several cellular processes such as sensing and migration [1]; they are also involved in cell-cell interactions [2, 45, 50, 51]. Consequently, filopodia have a very rich phenomenology, with lengths, growth and retraction rates, and lifetimes, highly variable. The mechanisms underlying formation, maintenance and dynamics of filopodia are complex and not yet fully understood. To shed some light on the growth dynamics of filopodia, we proposed a cross-talk between a theoretical stochastic model and experiments in PC3 cells.

We studied filopodia patterns in non-confluent PC3 cell cultures using confocal microscopy. Cells were fixed and inmunolabelled with rhodamine-phalloidin, allowing the visualization of actin structures. In some cases filopodia take the form of cell-cell bridges [3, 47, 51] implicated in the transport and interchange of molecules or vesicles. These structures are very stable and it has been suggested that they are reminiscent structures that remained after the retraction of the lamelipodia of two adjacent cells that were previously in contact [47]. We did not consider these kind of filopodial structures in this work. Rather, we focused on cell regions where single filopodia could be univocally tracked. We measured the lengths of these filopodia from the images using custom made tracking routines and determined their distribution.

Our analysis of the experimental results suggested the presence of two populations of cell regions according to the dispersion of their filopodial lengths, which we called Group 1 and Group 2. While Group 1 cells had filopodia with a narrow distribution of lengths, Group 2 showed a much more variable pattern.

These results were interpreted in terms of a stochastic model of filopodia growth that considers explicitly the main chemical reactions at the filament barbed end: polymerization and depolymerization of actin, as well as binding and unbinding of the regulatory proteins capping and formin. Also, the processes that result in the centripetal movement of actin filaments are represented in the model by an effective retrograde velocity. We inspected the effect of varying the retrograde velocity on the filopodial length and we found an inverse nonlinear dependence between them.

Considering these results, we proposed that the experimental lengths obtained for PC3 cells could be approximated to a linear combination of the *in silico* results for different values of retrograde velocities. In order to assess the reconstructed distribution, we compared its first 4 moments with the corresponding experimental ones. We found a good agreement between the model and the experimental data. We concluded that the reconstruction of Group 2 data demanded a wider range of values of *v*_*r*_ than Group 1 reconstruction, for which also low values of *v*_*r*_ were absent.

Based on these results, we proposed that Group 2 data came from cell regions where heterogeneous signaling induced further variability in the retrograde flow velocity or filopodia stability [21, 23–25] resulting in long filopodia and a wide length distribution. These cues could stem from the previous interactions with cells that were close before [52], or differential substrate adherence [53–55]. We did not consider explicitly these signaling processes and their effects in our model. Nevertheless, we included their effective behavior in the value of the retrograde flow velocity.

Although we cannot rule out other sources of variability in the lengths of filopodia, we believe that regulation of the retrograde flow represents an interesting mechanism by which cells can control other processes such as migration and/or invasion in the case of cancer cells.

## 4 MATERIALS AND METHODS

### 4.1 Numerical Simulations

We implemented an off-lattice one dimensional stochastic model of filopodial growth. We model growth dynamics of already initiated protrusions. A simulated filopodium consists of several actin filaments, in our model the maximum number of filaments contained by a filopodial bundle is 16. The filaments double helical conformation is simplified as single one dimensional structures. Each filament is an assembly of sub-units of size *δ* and is considered to be independent of the other filaments within the protrusion. As the computational model is one-dimensional we do not consider filaments radial distribution.

To simulate reactions and diffusion we apply a stochastic molecular approach. Results obtained with this method are expected to be equivalent to the ones found with the Gillespie algorithm, as shown by Erban at al. for a simpler model [31]. The model implemented here is similar to the proposed by Zhuravlev and Papoian [30]. It takes into account the following main components: (1) diffusion of molecules from the cell body into the filopodium compartment, (2) polymerization and depolymerization, (3) actin-binding regulatory proteins and (4) retrograde flow.

Let us consider a filopodium at time *t* composed of *N* (*t*) actin filaments of length *h*_*i*_(*t*), *i* = 1, 2, …, *N* (*t*). We define the filopodial length *H*(*t*) = *max*(*h*_*i*_) and consider the membrane position to be located *L/*2 = 25 nm above the longest filament *M* (*t*) = *H*(*t*) + *L/*2. A simulation time-step can be summarized as follows: (i) actin, formin and capping molecules can diffuse from the cytosol to the filopodia; (ii) all G-actin molecules whose position are between [*h*_*i*_(*t*) − *L*/2, *h*_*i*_(*t*) + *L/*2] have a non-zero probability of polymerizing at the *ith* filament, with on-rate values according to the presence of a formin and/or a capping molecule at the barbed end. For each actin polymerized, the filament length is increased by *δ*; (iii) each of the actin molecules at the barbed end can depolymerize; (iv) all the formin/capping molecules whose position are between [*h*_*i*_(*t*) − *L*/2, *h*_*i*_(*t*) + *L*/2] can bind into the barbed end of the *ith* filament (in the case that a formin/capping protein is not yet bound). Just one formin/capping can bind to the filament; (v) If there is a formin/capping bound to the filament, it can unbind; (vi) the filament length is reduced by the retrograde flow with a constant velocity *v*_*r*_. Steps (ii)-(vi) are performed for each of the *N* (*t*) filaments. The filopodial length is updated taking the value of the longest filament.

The simulation time-step was chosen so that all the probabilities are significantly smaller than 1 to guarantee stochasticity (δ*t* = 10^−6^*s*) and the algorithm was implemented in C programming language.

#### Diffusion

The diffusive molecules contemplated by our model are G-actin, capping protein and formin. The three types of biomolecules are present in the cytoplasm with different cytosolic concentrations (see Table 1), they follow Brownian motion dynamics and enter into the filopodial tube stochastically. G-actin, formin and capping protein diffusion into and within the filopodium is implemented in the same way Erban et al. did for G-actin [31] as we describe below.

G-actin is introduced into the filopodial structure with probability 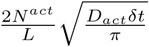, where *N*^*act*^ = 5.3 is the number of G-actin molecules within the volume *V* of a cylinder of height *L* = 50*nm* and diameter *d* = 150*nm* and is calculated by multiplying the bulk concentration [*A*]_*cyt*_ by *V*. The initial position of a molecule introduced into the filopodium is sampled from the probability density function 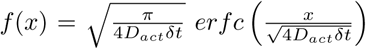 [31]. Formin and capping protein diffusive jumps into the filopodium are implemented likewise with the corresponding parameters (see Table 1), in these cases *N*^*form*^ = 0.042 and *N*^*cap*^ = 0.026. Flegg et al. [56] provide a detailed description of the implementation of diffusive jumps from a bulk domain (cytosol) into a molecular domain (filopodium).

All the molecules trajectories are explicitly computed. At time *t*, within the filopodium there are *n*^*act*^(*t*) G-actin molecules at positions 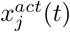, *j* = 1,2, …, *n*^*act*^(*t*), *n*^*cap*^(*t*) capping proteins located at 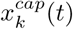, *k* = 1, 2, …, *n*^*cap*^ (*t*) and *n*^*form*^(*t*) formin molecules placed at 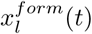, *l* = 1, 2, …, *n*^*form*^(*t*). The position of each particle can take values in a continuous range between the filopodial base and the membrane position [0, *M* (*t*)].

All the molecules positions evolve as random walk particles, hence an *mth* molecule located in *x*_*m*_(*t*) at time *t* moves to the position *x*_*i*_ (*t* + *δt*) = *x*_*m*_(*t*) + *δx*_*m*_, where the spacial step *δx*_*m*_ = *RMS* ⋅ *ϵ*_*m*_, RMS is the root mean square displacement in 1D 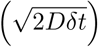 and *ϵ*_*m*_ is a standard normal distributed random number. Reflective boundary conditions were imposed at the membrane, if *x*_*m*_(*t*) + *δ*_*m*_ > *M*(*t*) then *x*_*m*_(*t* + *δt*) = *M* (*t*) − (*x*_*m*_(*t*) + *δx*_*m*_ − *M* (*t*)) and open boundary conditions were set at the filopodial base, if *x*_*m*_(*t* + *δt*) < 0 then the particle is removed from the system and the number of that type of molecules within the filopodial structure is reduced by one.

#### (De) Polymerization

At time *t* a filopodium consists of *N* (*t*) filaments of length *h*_*i*_(*t*), *i* = 1, 2, …, *N* (*t*). Each one of the filaments evolves with its own independent dynamics. Since we perform stochastic molecular simulations [31], for all the filaments we consider that each individual G-actin molecule located at a distance smaller than *L*/2 from the filament tip [29] can polymerize into its barbed end. In order to determine the assembling probability of each actin monomer, we must consider the rate of G-actin polymerization, 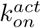, in units of [*s*^−1^] instead of [*μM*^−1^*s*^−1^] as usually reported (see Table 1). Therefore we take the volume of a cylinder of height *L* and diameter *d* and we get the parameter in the desired units 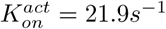 per molecule of G-actin. Accordingly, the assembling probability for each G-actin molecule within a time interval (*δt*) is 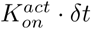. The case where there is a formin and/or a capping molecule bound to the filament will be discussed next.

Because we model the microfilaments as single one-dimensional objects, while they actually are double helices, the filaments increase its lengths by half the size of an actin monomer (*δ*) when polymerization occurs. Next, we remove the polymerized G-actin protein from the pool of free molecules. Moreover, within a time interval (*δt*) each filament has a depolymerizing probability of 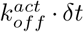, as 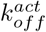 is the dissociation rate of an actin monomer at the barbed end. If a depolymerization event happens the filament length is reduced by *δ* and a new G-actin monomer is introduced at the position of the filament length before depolymerizing *h*_*i*_(*t*).

#### Actin-binding regulatory proteins

Two types of actin-binding proteins are contemplated by the model: formin and capping proteins. In the same way that polymerization was implemented, for each actin filament all the formin and capping molecules located at a distance less than *L*/2 can attach to the filament with the appropriate probability. In order to calculate these probabilities, the units of the on-rates were transformed from [*μM*^−1^*s*^−1^] to [*s*^−1^], as it was done with the polymerization rate 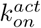. Thereby we obtain 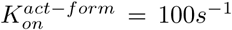, 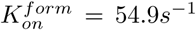, 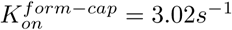, 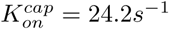 and 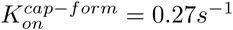.

Formin (un)binds to the free filaments barbed ends with probability 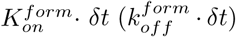. When a filament is anticapped with formin the polymerization probability is 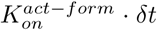 whilst depolymerization remains unaltered. On the other hand, capping proteins can bind to each one of the filaments free barbed ends with probability 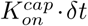 completely blocking polymerization and depolymerization. Uncapping of capped barbed ends takes place with probability 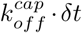. A capping protein can bind to a formin-bound barbed end with probability 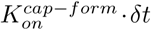 and unbind with probability 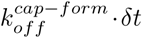 as well as formin can (un)bind to a capped barbed end with probability 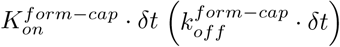. In analogy with the G-actin dynamics implementation, when a formin or a capper binds to a filament barbed end the protein is removed from its respective pool and the number of the corresponding type of molecule is reduced by one. In addition, when unbinding occurs the new unbound protein is introduced into the system of free molecules in the same way that we did for G-actin when a depolymerization event took place.

#### Retrograde flow

Microfilaments pointed end depolymerization together with the action of myosin leads to the phenomenon of retrograde flow, a centripetal movement of actin filaments. Our model assumes all the filaments that conform a filopodial bundle move backwards with the same constant retrograde velocity *v*_*r*_. At every time-step *δt* each filament of length *h*_*i*_(*t*) is shortened by *v*_*r*_ ⋅ *δt* therefore *h*_*i*_(*t* + *δt*) → *h*_*i*_(*t*) − *v*_*r*_ ⋅ *δt*.

### 4.2 Statistical analysis

The filopodial length averaged over an ensemble of *n* simulation runs at time *t* is defined as:

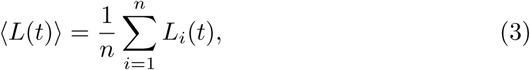

where *L*_*i*_(*t*) is the length of the realization *i* at time *t*. Obviously, the *n* realizations refer to the same parameters values. Further, the mean filopodial length is defined as the time average of 〈*L*(*t*)〉 over *T* values:

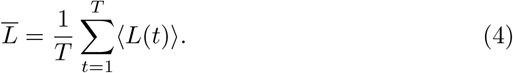

Since we choose a sampling time of 1s, *T* is the maximum time considered (in seconds).

In order to calculate 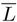 (shown in Fig. 2b) we consider *n* different runs of *T* = 1000*s*, with a time step of 1s. For *v*_*r*_ between 20 nm/s and 100 nm/s we use *n* = 5, whereas for *v*_*r*_ = 5 and *v*_*r*_ = 10 we take *n* = 18 and 17, respectively. Better statistics are necessary to represent low *v*_*r*_ data because of large fluctuations in the transient period. The same simulation data were used to calculate the probability density functions *S*_*j*_ (Step 2 of Section 4.5). In cases where binning was required, the size of the bins was determined by Freedman–Diaconi’s rule. Boxplots are presented as mean, SEM and SD, according to [49]. For the filopodial lengths analysis, distributions were compared with a two-sample Kolmogorov–Smirnov test. A kernel density estimation was applied to the data in order to obtain probability density estimate curves.

### 4.3 Cell culture, samples preparation for imaging and confocal microscopy

PC3 cells were obtained from the American Type Culture Collection (Manassas, VA, USA) and were routinely cultured in RPMI 1640 (Invitrogen, Grand Island, NY, USA) supplemented with 10% fetal bovine serum (FBS), penicillin 100 U/mL, streptomycin 100 g/mL and amphotericin 0.5 g/mL. The rhodamine–phalloidin was purchased from Life Technologies (Thermo Fisher Scientific Inc., Eugene, OR, USA). Immunofluorescence experiments and quantitative microscopy: PC3 cells were fixed with 8% paraformaldehyde (PFA) (20 min, room temperature) and stained with rhodamine–phalloidin (1 h, room temperature) [45]. Confocal images were acquired by confocal microscopy (FV1000, Olympus, Tokyo, Japan) using an UPlanSApo 60x oil immersion objective (NA 1/41.35; Olympus), a diode laser of 543 nm as the excitation source and fluorescence was collected in the range of 555–655 nm.

### 4.4 Filopodia localization and tracking

Image processing was done with ImageJ and data analysis was carried out with R. The workflow to determine filopodial length involved manual tracking of filopodia and the latter automatic determination of its length. Image contrast and brightness were adjusted for better identification of filopodia. Protrusions that did not accomplish certain minimal requirements were not measured as stated before.

We acquired intensity profiles of lines traced above the filopodia using the ImageJ line profile tool in segmented mode, line width was established considering image resolution so that each filopodium was completely covered by the line (7 pixels for 1 image, 8 pixels for 8 images and 10 pixels for the remaining 39).

The program computes the average intensity along the line. The line starting point was set inside the cell body and the ending point was placed beyond the visible tip as shown in Fig.3e. We also acquired background intensity profiles for every image. Each filopodium and cell was assigned an ID number, and for every cell we also kept information if it had contacts or not with surrounding cells. All the objects were saved within the images so that every filopodium can be tracked back.

The recovered intensity profiles showed a rapid rising until reaching a maximum around the filopodia base and the subsequent gradual decrease until reaching the filopodia extreme where the signal becomes indistinguishable from the statistical background noise (Fig.3f). Then, each intensity profile was smoothed with the R smooth function, a median filter to reduce the noise signal. We set the pixel with the highest intensity as base of the filopodia. We performed statistical analysis on the background intensity profiles for every image, determining the mean background signal and its standard deviation. The tip of the structure was determined by setting a threshold as the mean plus 3 standard deviations of the background signal. We obtained filopodial length subtracting the tip to the base position.

We aimed to estimate the sampled filopodia percentage from the total population within the explored regions. For that, we measured the contour length of 45 sampled areas (≈ 1194 *µm*) and multiplied it by the filopodial linear density (≈ 0.5 *µm*^−1^) [45]. This led to ≈ 597 expected filopodia in contrast to 337 ones within the covered perimeter. Therefore, we estimated to have measured the length of about ≈56% of the protrusions within the explored areas.

### 4.5 Reconstruction of the experimental filopodium length distributions

As we stated in Section 2.4, the probability density function of the filopodium length for Groups 1 and 2 obtained from experimental data can be reconstructed by a linear combination of the simulated distributions obtained for different values of retrograde velocity *v*_*r*_. In order to determine the weight of each simulation distribution in the reconstruction of the experimental data, the procedures are detailed below.

**Step 1:** We use the kernel density estimation to generate two distributions, *E*_1_ and *E*_2_, for the experimental filopodium lengths for Groups 1 and 2, respectively.

**Step 2:** We use the kernel density estimation to generate *m* length distributions from the simulation data, each one corresponding to a different value of *v*_*r*_. Let us call these distributions as *S*_*j*_, with *j* = 1,…, *m*. For the simulations we consider the following *m* = 11 values of retrograde velocity, *v*_*r*_ = 5, 10, 20, 30, 40, 50, 60, 70, 80, 90, 100 nm/s. Information about the simulation data considered to construct the functions *S*_*j*_ are detailed in Section 4.2.

**Step3:** To reconstruct the experimental distributions *E*_1_ and *E*_2_, we should determine the weights 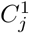 and 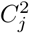 of each simulation distribution *S*_*j*_, for Groups 1 and 2, respectively. That is: 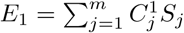 and 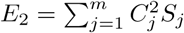 (Eqns. 1 and 2, respectively).

**Step 4:** The moments of order *i*, *µ*_*i*_, with *i* = 1,…, 4 are calculated for the experimental distributions *E*_1_ and *E*_2_ (see Table 4). Therefore, we can define 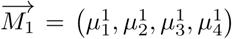 and 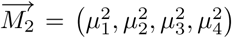, for Groups 1 and 2, respectively.

**Table 4.**
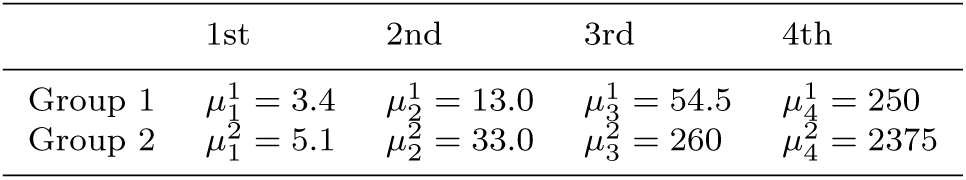
Moments of the experimental distributions, *E*_1_ and *E*_2_.

|  | 1st | 2nd | 3rd | 4th |
| --- | --- | --- | --- | --- |
| Group 1 | $\mu_1^1 = 3.4$ | $\mu_2^1 = 13.0$ | $\mu_3^1 = 54.5$ | $\mu_4^1 = 250$ |
| Group 2 | $\mu_1^2 = 5.1$ | $\mu_2^2 = 33.0$ | $\mu_3^2 = 260$ | $\mu_4^2 = 2375$ |

**Step 5:** We compute the moments of order *i*, *µ*_*i*_, with *i* = 1,…, 4, for each of the *S*_*j*_ simulated data performed for different retrograde velocities values: 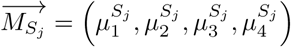, with *j* = 1,…, *m*.

**Step6:** We consider the following linear regression for each of the moments of Group 1:

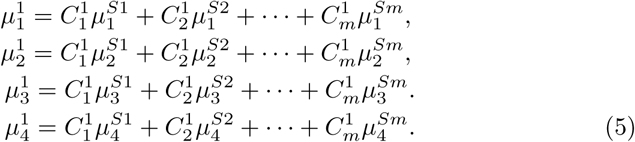

Similar equations are found for Group 2.

**Step 7:** Let us normalize the moments of the experimental distributions respect to the first moment as:

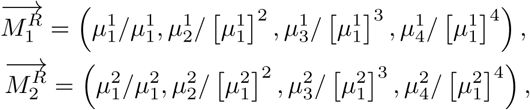

Therefore, Eqs. 5 for Group 1 and similar equations for Group 2 can be rewrite (in a reduced way) as:

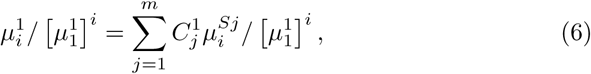

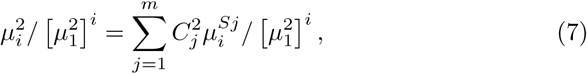

with *i* = 1,…, 4.

This normalization procedure is necessary in order to give similar importance to the different moments when the regression analysis is done. In other way, high moments will be favored.

**Step8:** We determine the coefficients 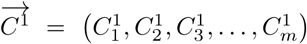 and 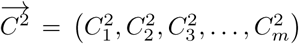, associated with the constrains given by Eqs. 6 and 7, respectively. The system has no exact solution, therefore we use a L1 minimization [57] to obtain 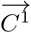 and 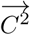.

**Step9:** After the determination of 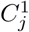 and 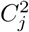 we normalize the coefficients 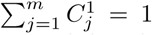 and 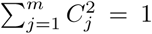 to ensure the normalization of the reconstructed distributions.

